# Possible Novel Sulfolipid Utilization Pathway in Giant Clams and Other Aquatic Invertebrates: Implications for Photosymbiosis and Sulfur Cycling

**DOI:** 10.64898/2026.02.11.705467

**Authors:** Taiga Uchida, Hiroshi Yamashita, Go Shimada, Eiichi Shoguchi, Chuya Shinzato

## Abstract

Photosymbiosis with dinoflagellates of the family Symbiodiniaceae enables giant clams to thrive in oligotrophic coral reef environments. However, mechanisms by which clams utilize algal-derived biomolecules remain largely unexplored. Using newly available genome resources for photosymbiotic bivalves (*Tridacna* and *Fragum*), we conducted a comparative genomic analysis to identify positively selected genes from these photosymbiotic bivalve lineages, that are potentially involved in symbiotic adaptations. Among candidate genes, we focused on sulfoquinovosidase (SQase), an enzyme that hydrolyzes sulfoquinovose (SQ) from sulfoquinovosyl diacylglycerol (SQDG), a sulfur-containing sulfolipid abundant in photosynthetic membranes. Although SQDG degradation has been characterized in bacteria, the distribution and role of SQase in animals have not been systematically examined. We found that possibly functional SQase homologs are widely distributed among aquatic invertebrates, but are largely absent in terrestrial taxa. In silico predictions showed that most animal SQases possess signal peptides and enter the secretory pathway, with lineage-specific gains and losses of membrane association, suggesting functional diversification. Transcriptomic analyses further demonstrated that SQase is predominantly expressed in digestive organs in diverse taxa, whereas in giant clams, it is also highly expressed in the outer mantle, the tissue primarily involved in harboring symbionts enriched in SQDG. Together, these results suggest that in aquatic invertebrates, SQase functions as a digestive enzyme for algal-derived sulfolipids and that photosymbiotic bivalves have expanded its deployment to tissues specialized for symbiosis. Our findings uncover a previously unrecognized metabolic interface between photosymbiosis and sulfur utilization in aquatic invertebrates, with broader implications for host-symbiont interactions and marine sulfur cycling.

## Introduction

Symbioses between animals and microorganisms are widespread in nature and are thought to have driven acquisition of novel traits, environmental adaptation, and diversification throughout evolution (Gilbert et al. 2012; McFall-Ngai et al. 2013; Melo Clavijo et al. 2018). Among these, photosymbiosis, the association between photosynthetic organisms and heterotrophic hosts, confers substantial advantages for adaptation to oligotrophic marine environments. In coral reef ecosystems, dinoflagellates of the Family Symbiodiniaceae form symbioses with reef-building corals and giant clams, supplying photosynthetically fixed carbon to their hosts and thereby sustaining the high productivity of these ecosystems (Baker 2003; Roth 2014; Blackall et al. 2015; Mies 2019). In return, host organisms provide inorganic nutrients such as carbon, nitrogen, and phosphate, as well as a protected microhabitat, to their symbiotic dinoflagellates (Yellowlees et al. 2008).

Photosymbiosis with dinoflagellates of the Family Symbiodiniaceae occurs in a broad range of taxa, including animals such as cnidarians, molluscs, sponges, and xenacoelomorphs, as well as protists such as foraminiferans and ciliates. Photosymbiosis is thought to have evolved independently in each host lineage (Mies et al. 2017; Melo Clavijo et al. 2018). Among molluscs, the most representative photosymbiotic taxa are giant clams (genera *Tridacna* and *Hippopus*, subfamily Tridacninae, family Cardiidae), which have served as model systems for recent studies on mechanisms and evolution of molluscan photosymbiosis (Armstrong et al. 2018; Mies 2019; Ip & Chew 2021; Li et al. 2025). Giant clams are bivalves that inhabit tropical and subtropical coral reefs in the Indo-Pacific Ocean. They are well known for their exceptionally large body sizes and vividly colored mantles (Mies 2019). Their symbiotic dinoflagellates reside extracellularly in specialized tubular structures, termed “zooxanthellal tubes”, that extend from the stomach to and throughout the outer mantle (Norton et al. 1992). Classical histological observations suggested that in the outer mantle, amoebocytes selectively engulf a subset of symbiotic dinoflagellates (Morton 1978), and that intracellular digestion by amoebocytes may occur in the hepatopancreas (digestive diverticula) (Fankboner 1971). However, more recent studies have demonstrated that large numbers of dinoflagellate cells are present in fecal pellets of giant clams (Morishima et al. 2019), indicating that some symbionts may be transported directly through the digestive tract and potentially subjected to extracellular degradation.

Another lineage of photosymbiotic bivalves includes heart cockles. In the Family Cardiidae, a monophyletic group of species belonging to the subfamily Fraginae (genera *Fragum*, *Corculum*, and *Lunulicardia*) harbors symbiotic dinoflagellates (Kirkendale 2009). The subfamilies Tridacninae and Fraginae are thought to have acquired photosymbiosis independently (Li et al. 2020). As in clams, heart cockles host dinoflagellates primarily in tubular structures in the mantle; however, their mantles are considerably less developed than those of giant clams (Farmer et al. 2001; Kirkendale 2009; Li et al. 2018).

When organisms acquire novel traits and adapt to new environments, changes in protein function driven by amino acid substitutions can prove essential. In reef-building corals, several studies have explored genes exhibiting elevated rates of evolution or signatures of positive selection relative to membrane transport, stress responses, and pattern recognition, with possible links to symbiosis (Voolstra et al. 2011; Iguchi et al. 2011; Shinzato et al. 2021). In these symbionts, genes associated with photosynthesis, membrane transport, and starch biosynthesis are under positive selection in dinoflagellate lineages that engage in symbiosis with animals, suggesting their involvement in symbiosis (Liu et al. 2018; Ishii et al. 2025). In contrast, comparative genomic and molecular evolutionary analyses have been largely lacking in photosynthetic bivalves, partly due to the limited availability of genomic resources. Recently, however, genome sequences and associated analyses of giant clams have been reported by several groups (Jun Li et al. 2024; Li et al. 2025; Uchida, Yamashita, et al. 2025), and multiple genomes of *Tridacna* and *Fragum* have been released through the Aquatic Symbiosis Genomics project led by the Wellcome Sanger Institute (McKenna et al. 2021). These advances have now made photosymbiotic bivalves amenable to comparative genomic analyses.

In this study, we leveraged recently sequenced genomes of *Tridacna* and *Fragum* to identify candidate genes that have experienced positive selection in these lineages, and that may be involved in symbiotic adaptations. Among these candidates, we focused on a gene encoding sulfoquinovosidase (SQase). SQase catalyzes the hydrolytic release of sulfoquinovose (SQ) from sulfoquinovosyl diacylglycerol (SQDG), a sulfur-containing sulfolipid abundant in chloroplasts of photosynthetic organisms, and sulfoquinovosyl glycerol (SQGro), which is generated by deacylation of SQDG. Thus, SQase represents the gateway enzyme of the SQDG degradation (sulfoglycolytic) pathway (Speciale et al. 2016). SQ is one of the most abundant organic sulfur compounds on earth, and both the biosynthesis and degradation of SQDG are critical in global sulfur and carbon cycling (Goddard-Borger & Williams 2017; Snow et al. 2021). Although mammals can deacylate SQDG with lipases (Gupta & Sastry 1987), there has been no report that they can liberate SQ. Instead, at least in mammals, this step is thought to be mediated by gut bacteria possessing SQase genes (Hanson et al. 2021; Krasenbrink et al. 2025). Here, we investigated the distribution of SQase homologs throughout the animal kingdom and inferred their molecular functions and subcellular localization based on amino acid sequence analyses. We further characterized their gene expression patterns using RNA-seq data. Our results provide new insights into mechanisms related to photosymbiosis in giant clams and reveal a previously underappreciated metabolic pathway in aquatic invertebrates, with broad implications for sulfur and carbon cycling in marine ecosystems.

## Results

### Identification of Candidate Genes for Positive Selection in Photosymbiotic Bivalves

The number of single-copy orthologs shared by 12 bivalve species was 2,014. The maximum likelihood molecular phylogenetic tree constructed using concatenated amino acid sequences of these orthologs supported the monophyly of *Tridacna* and *Fragum* (Fig. 1a). Using this species phylogeny as the guide tree, 2014 single-copy orthologs were subjected to a survey of positive selection on the foreground branch leading to the exclusive common ancestor of *Tridacna* and *Fragum*, with CODEML (Yang 2007). Numbers of candidate genes for positive selection (likelihood ratio test *p*-value < 0.05) were 175 and 360 for *Tridacna* and *Fragum*, respectively. Among these genes, 54 were shared by both genera (Fig. 1b). Shared candidates included genes encoding putative NPC intracellular cholesterol transporter 1 (NPC1), sulfoquinovosidase (SQase), and monocarboxylate transporter (SLC16). *Tridacna*-specific candidates included genes for sodium-independent sulfate anion transporter (SLC26A11), solute carrier family 15 member 4 (amino acid and oligopeptide transporter, SLC15A3), sodium-dependent glucose transporter, GQ-rhodopsin (Fig. 1b). Full results are shown in Tables S1 and S2.

**Fig. 1.**
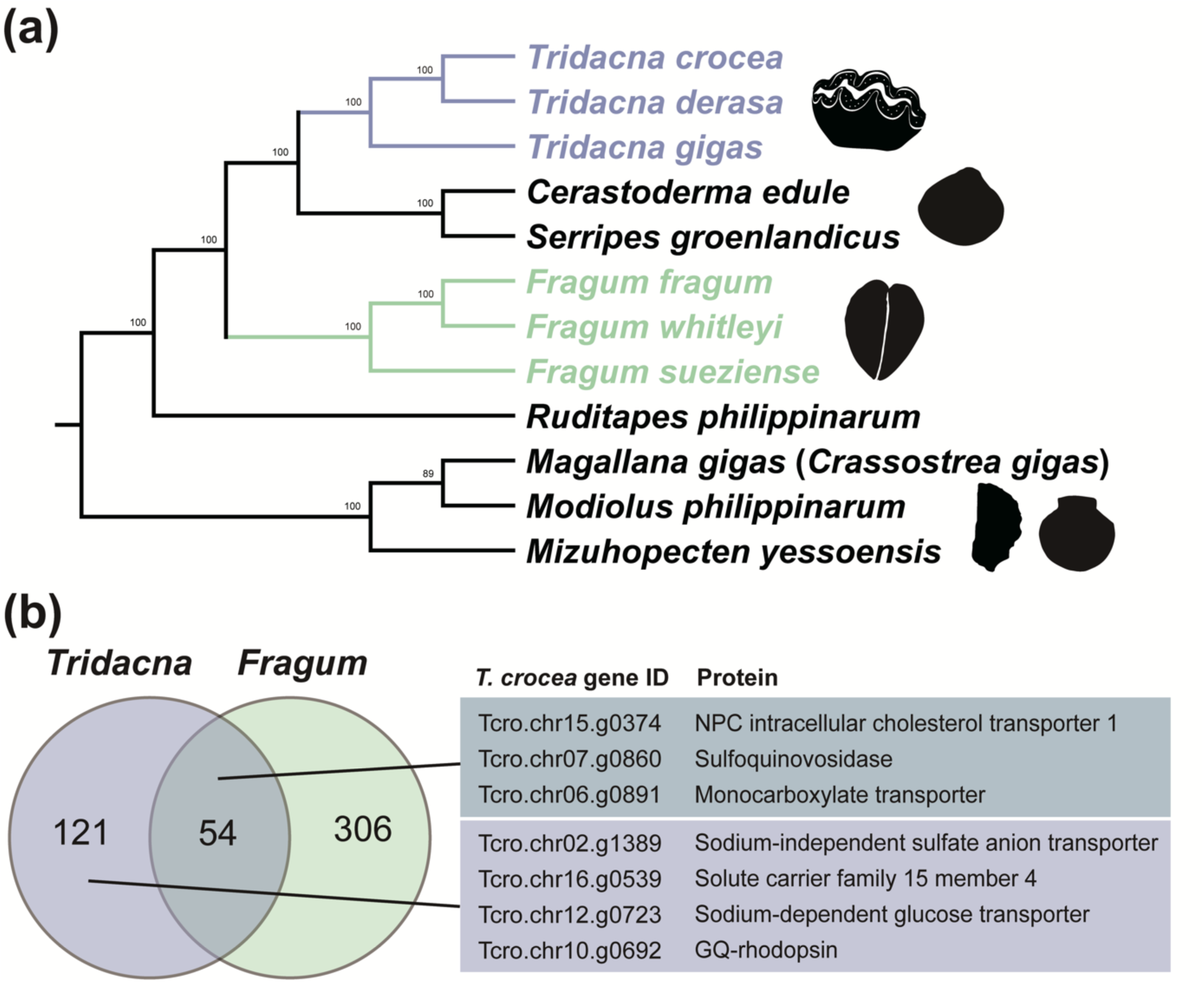
Overview of analysis for positive selection. **a)** Phylogenetic relationships of 12 bivalve species used for detecting positive selection. A maximum likelihood molecular phylogenetic tree was constructed using concatenated amino acid sequences of 2,014 single-copy orthologs shared among these species. Bootstrap values are shown on branches. Silhouette images of *Tridacna* and *Magallana* were downloaded from PhyloPic. **b)** A Venn diagram of candidate genes for positive selection in each lineage (left) and representative genes of particular interest (right). The full list of candidate genes is shown in Tables S1 and S2.

### Distribution of Sulfoquinovosidase (SQase) Homologs among Animals

Among candidate genes, we focused on one encoding a protein with sequence similarity to sulfoquinovosidase (SQase). SQase is generally known as a bacterial protein (Snow et al. 2021), not as an animal protein; therefore, we explored SQase homologs among animals. Three-step exploration of SQase homologs suggested conservation of SQase homologs among various lineages of animals, including the Arthropoda, Mollusca, Annelida, Brachiopoda, Bryozoa, Hemichordata, Echinodermata, Chordata, and Rotifera (Figs. 2, 3, and Figs. S1-3). In the maximum likelihood phylogenetic tree constructed using known glycoside hydrolases as outgroups, topologies of SQase candidate sequences, except for those from the Rotifera, were generally consistent with generally accepted phylogenetic relationships among corresponding taxa, although bootstrap support values were moderately low overall. In contrast, only sequences derived from Rotifera formed a clade with bacterial SQases, supported by a bootstrap value of 74%.

**Fig. 2.**
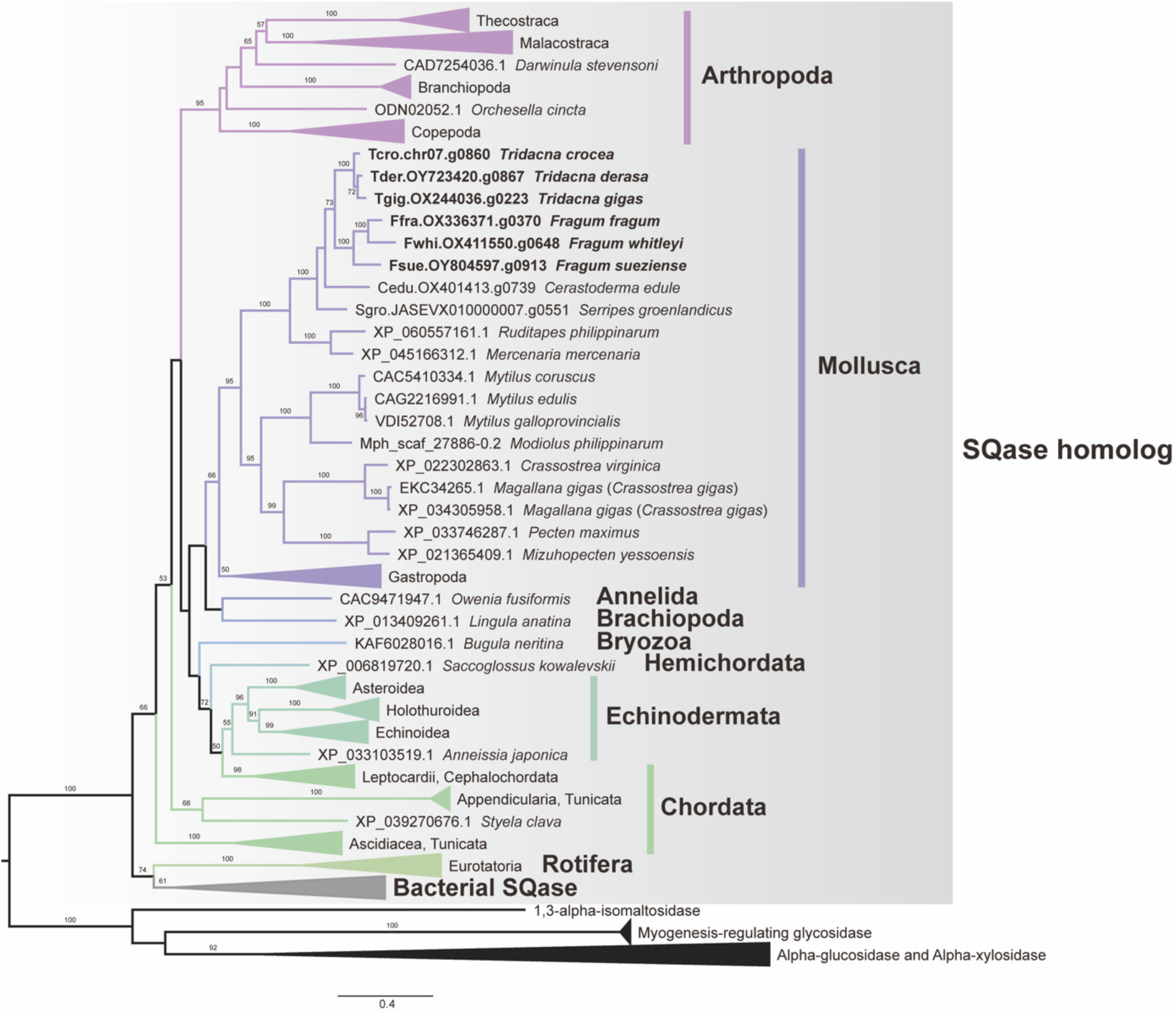
Phylogenetic relationships of animal SQases and bacterial homologs. Bootstrap values > 50% are shown on branches.

**Fig. 3.**
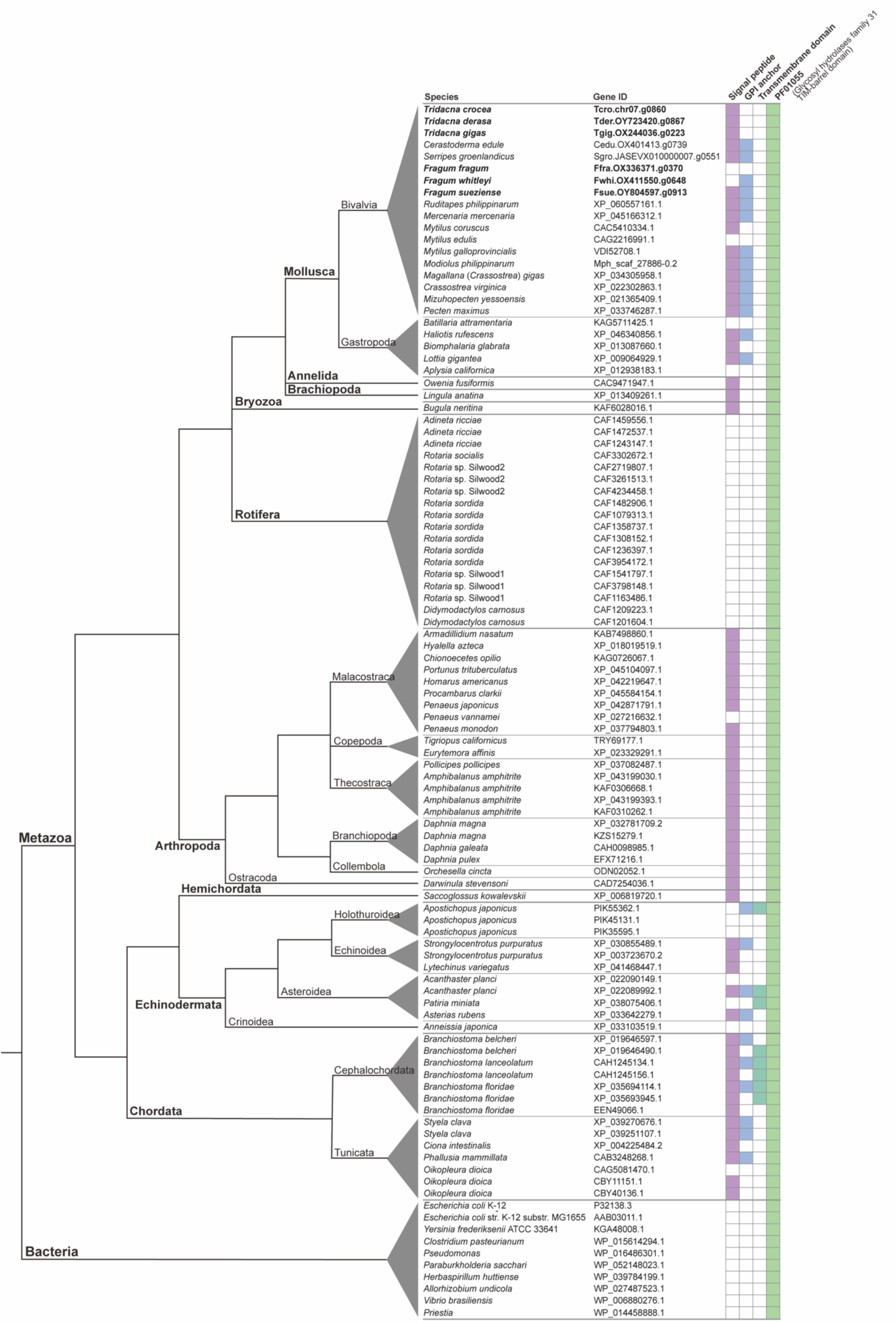
Predicted domain structure of animal SQases and bacterial orthologs. The cladogram on the left represents species phylogeny based on previous studies (Dunn et al. 2014; Giribet & Edgecombe 2019; Rahman & Zamora 2024). For each protein, presence or absence of a predicted N-terminal signal peptide, GPI-anchor attachment site, transmembrane domain, and glycosyl hydrolase family 31 TIM-barrel domain (PF01055) is indicated by the grid on the right. Both the transmembrane domain (when present) and the PF01055 domain were detected as single copies in all proteins shown.

In total, SQase homologs were identified in 65 animal species in this study. Among them, only two terrestrial species, *Armadillidium nasatum* (Malacostraca, Arthropoda; woodlouse) and *Orchesella cincta* (Hexapoda, Arthropoda; springtail), were detected, whereas all remaining species were aquatic invertebrates (Figs. 2 and 3). In the Mollusca, SQase homologs were detected in bivalves and gastropods, but not in cephalopods. In the Annelida, SQase homologs were identified only in *Owenia*, which represents one of the earliest-branching annelid lineages.

To investigate the presence of genes encoding the first enzymes of SQ degradation pathways downstream of SQase, TBLASTN homology searches were performed against both gene models and the genome sequence of *T. crocea*. In the search of gene models, several sequences showed similarity to *Pseudomonas putida* sulfoquinovose 1-dehydrogenase (UniProt accession: P0DOV5). However, all of these sequences were annotated as dehydrogenases/reductases, distinct from sulfoquinovose 1-dehydrogenase, based on SwissProt BLAST annotations (Table S3). In the search against the genome sequence, all significant hits corresponded to protein-coding regions. With the exception of one gene annotated as an ADP-ribosylation factor, all other hits were annotated as dehydrogenases/reductases distinct from sulfoquinovose 1-dehydrogenase, based on SwissProt BLAST annotations (Table S4).

### Molecular Features of Animal SQase

To predict signal peptides that regulate extracellular protein targeting, the machine-learning-based tool SignalP 6.0 was used (Teufel et al. 2022). Signal peptides were predicted at the N-terminus of SQase proteins in many animal lineages, except for the Rotifera, in which no SPs were detected in any species examined (Figs. 3 and 4b). Prediction of glycosylphosphatidylinositol (GPI) anchor attachment using NetGPI 1.1 (Gíslason et al. 2021) suggested the presence of GPI-anchor signals in many molluscs other than *Tridacna*, and in some echinoderms and chordates. Transmembrane helices predicted by DeepTMHMM 1.0.44 (Hallgren et al. 2022) were detected only in some echinoderms and chordates, but not in molluscs. No signal peptides, GPI-anchor attachment sites, or transmembrane helices were predicted in bacterial or rotifer SQase sequences (Figs. 3 and 4b).

**Fig. 4.**
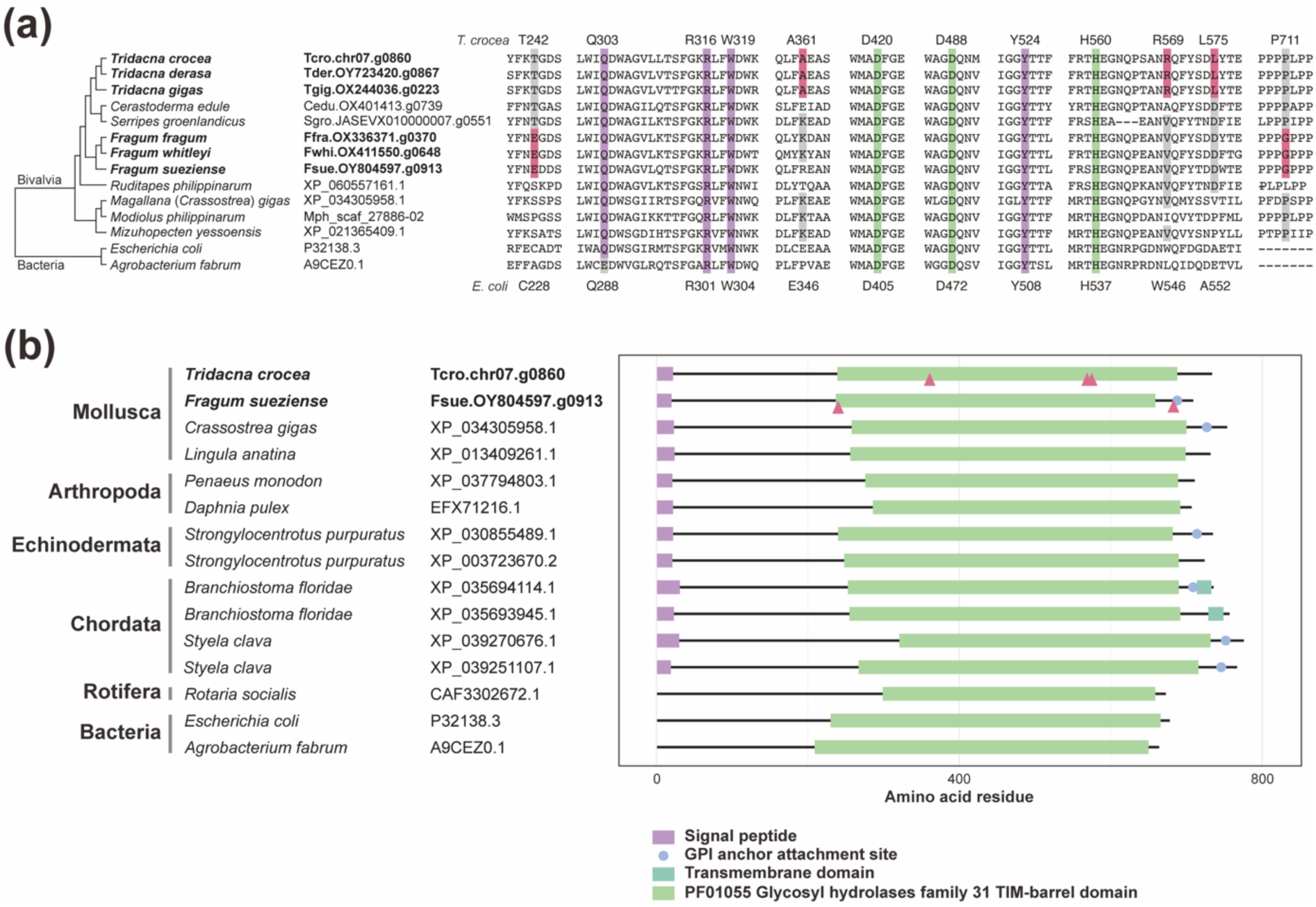
Important amino acids and predicted structure of animal SQases. **a)** Amino acid sequence alignment of animal SQases of representative species. Residue numbers of *T. crocea* animal SQase are shown on the top, and those of *E. coli* YihQ (SQase) are shown on the bottom. Substrate recognition in *E. coli* SQase relies on conserved amino acids, including Q288 for hydroxyl recognition and R301, W304, and Y508 for specific recognition of the sulfonate group of SQ. These amino acids are highlighted in purple. *E. coli* SQase D405 and D472 function as the catalytic nucleophile and acid-base catalyst, respectively. H537 also contributes to substrate binding. These amino acids are highlighted in green. In *A. fabrum* SQase, substitution of Q288 with glutamate is compensated by a paired substitution of Q262 with lysine (K245) (Speciale et al. 2016; Abayakoon et al. 2018). Amino acids under positive selection are highlighted in red. **b)** Diagram of the predicted domain structure and amino acids under positive selection.

Amino acids previously reported as critical for SQase activity and substrate specificity by Speciale et al. (2016) were well conserved in animal SQase homologs (Fig. 4a). Protein sequences in which multiple substitutions or deletions were observed at these seven key amino acids were excluded from the phylogenetic and molecular feature analyses shown in Figs. 2-4. Likewise, only sequences containing a single glycosyl hydrolase family 31 TIM-barrel domain (PF01055), as observed in known SQases, were reported as animal SQase homologs in this study (Figs. 3 and 4).

### Reassessment of Positive Selection in Bivalve SQase

As described above, signal peptides and GPI-anchor attachment were predicted in Animal SQase proteins. Because signal peptides and amino acids located downstream of GPI-anchor attachment sites (ω-sites) are removed during protein maturation, these regions are expected to be under relatively relaxed functional constraints. Consistently, multiple sequence alignments of animal SQases showed reduced alignment quality near both the N- and C-termini (data not shown). To minimize the influence of these regions on downstream analyses, they were removed, and only amino acid sequences likely retained in the mature protein were reanalyzed. A gene phylogeny reconstructed from the trimmed alignment exhibited overall lower bootstrap support and partial topological incongruence compared with species phylogeny; however, no major conflicts violating assumptions of subsequent analyses were observed (Fig. S4).

Using the species phylogeny as the guide tree, positive selection was detected on branches leading to *Tridacna* (*p* = 0.00268, dN/dS = 16.23) and *Fragum* (*p* = 0.000900, dN/dS = 999.00). Similar results were obtained when the gene phylogeny was used as the guide tree, with significant signals of positive selection detected for both *Tridacna* (*p* = 0.00135, dN/dS = 14.49) and *Fragum* (*p* = 0.00181, dN/dS = 999.00). In both analyses, three amino acids in *Tridacna* and two in *Fragum* were inferred to be under positive selection (Fig. 4a). All positively selected sites were located in or immediately downstream of the glycosyl hydrolase family 31 TIM-barrel domain and did not coincide with amino acids previously identified as critical for SQase activity or substrate specificity (Speciale et al. 2016).

### SQase Gene Expression Patterns in Several Aquatic Invertebrates

Tissue-specific expression patterns of animal SQase genes were compared using RNA-seq data obtained from previous studies and publicly available databases. In *T. crocea*, SQase gene expression levels were significantly higher in the hepatopancreas and outer mantle than in adductor muscle, gill, kidney, and pedal mantle (Fig. 5). Similarly, in *Tridacna squamosa*, SQase gene expression was high in the hepatopancreas and outer mantle, whereas expression levels in other tissues were extremely low.

**Fig. 5.**
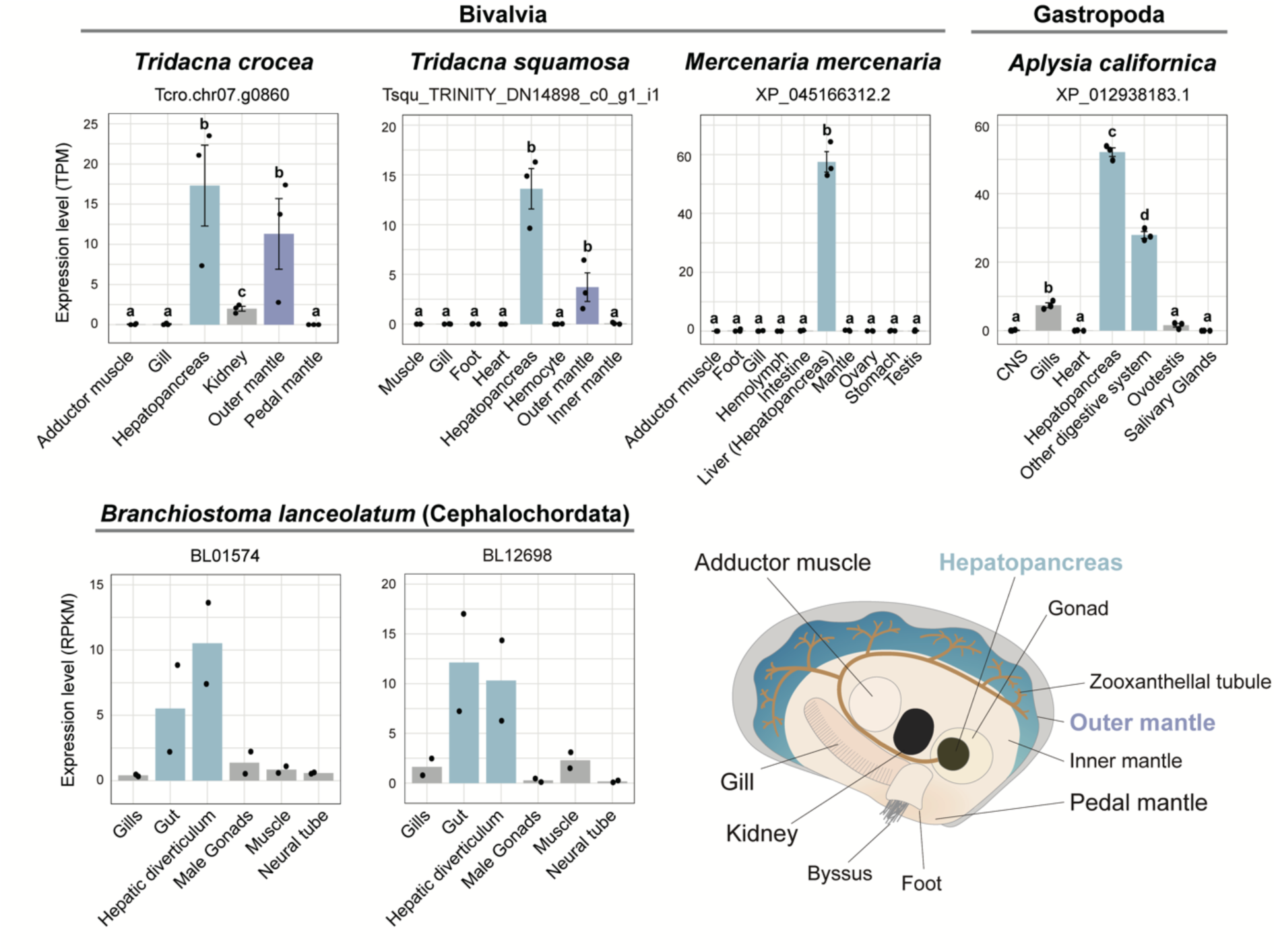
Tissue-specific gene expression of animal SQases. Expression levels of animal SQase genes based on publicly available RNA-seq data and a schematic diagram of the anatomy of *T. crocea* are shown. Digestive tissues are highlighted in blue, and the outer mantle is highlighted in purple. a-d represents statistically indistinguishable groups (*p* > 0.05).

In bivalve species other than giant clams, SQase expression was high in the liver (hepatopancreas) of *Mercenaria mercenaria*, whereas expression levels in other tissues were extremely low (Fig. 5). Among non-bivalve molluscs, *Aplysia californica* showed significantly higher SQase expression in the hepatopancreas than in six other tissues analyzed, followed by significantly higher expression in the other digestive system than in the remaining five tissues.

In invertebrates other than the Mollusca, SQase expression was examined in *Branchiostoma lanceolatum* (European lancelet). Although no statistical tests were performed due to the small sample size (n = 2), both individuals showed a tendency toward higher SQase expression in the gut and hepatic diverticulum, whereas expression levels in other tissues were low (Fig. 5).

## Discussion

### Lineage-Specific Positive Selection Possibly Associated with Photosymbiosis

Photosymbiosis between dinoflagellates of the Family Symbiodiniaceae and animals is thought to have been acquired independently multiple times during animal evolution, and is essential for sustaining primary production in coral reef ecosystems (Mies et al. 2017; Melo Clavijo et al. 2018; Li et al. 2020). Among bivalves, photosymbiosis with dinoflagellates occurs in two distinct lineages (Fig. 1a); however, little is known about genes that contributed to acquisition of this symbiotic lifestyle. In this study, we utilized recently available genomic resources for giant clams (*Tridacna*) and heart cockles (*Fragum*) (McKenna et al. 2021; Uchida, Yamashita, et al. 2025) to explore genes that have experienced positive selection in these lineages, and that may have been involved in development of symbiosis.

As a result of branch-site analyses using CODEML, we identified 175 and 360 candidate genes showing signatures of positive selection in *Tridacna* and *Fragum*, respectively. Candidate genes that were significant only in *Tridacna* included a sodium-independent sulfate anion transporter (SLC26A11), solute carrier family 15 member 4 (an amino acid and oligopeptide transporter, SLC15A3), a sodium-dependent glucose transporter, and GQ-rhodopsin (Fig. 1b). Among these, transporter genes may be involved in exchange of metabolites between giant clams and their symbionts. Indeed, in corals, expression of several transporter genes increases during establishment of symbiosis with dinoflagellates, and these genes have therefore been proposed as candidates involved in symbiosis (Yuyama et al. 2018; Yoshioka et al. 2023). In addition, in giant clams, members of the SLC26 family have been hypothesized to mediate the supply of bicarbonate (HCO₃⁻) or nitrate (NO₃⁻) to symbionts (Uchida, Yamashita, et al. 2025). Giant clams extend their mantles outside the shell to provide light for symbiont photosynthesis and rapidly move the mantle in response to changes in the light environment (Soo & Todd 2014); therefore, GQ-rhodopsin may be involved in such light-responsive behaviors. These genes are conserved in a wide range of bivalve lineages (Fig. 1a), and amino acid substitutions that have accumulated in proteins they encode may have contributed to adaptations specific to giant clams, including photosymbiosis.

We identified 54 candidate genes that were shared between *Tridacna* and *Fragum*. These included genes encoding NPC intracellular cholesterol transporter 1 (NPC1) and sulfoquinovosidase (SQase) (Fig. 1b). NPC1 functions together with NPC intracellular cholesterol transporter 2 (NPC2) in intracellular sterol transport (Subramanian & Balch 2008). In cnidarians such as corals and sea anemones, these proteins are thought to contribute to symbiosis through acquisition of symbiont-derived sterols (Dani et al. 2014, 2017; Hambleton et al. 2019; Levy et al. 2021; Yoshioka et al. 2023; Shikina et al. 2024). In *Fragum*, NPC2 gene expression in the mantle increases in response to light, whereas such patterns were not observed in the foot of *Fragum* or in closely related non-symbiotic species (Ruiqi Li et al. 2024). Furthermore, in *T. crocea*, several NPC2 homologs are expressed specifically and at extremely high levels in the outer mantle, and their expression is downregulated following breakdown of symbiosis (Uchida, Yamashita, et al. 2025). These results suggest possible involvement of NPC2 in photosymbiosis in bivalves. In these studies, NPC1 did not exhibit tissue-specific expression or expression changes associated with light or symbiosis breakdown; however, it is possible that positive selection has led to functional changes in NPC1 that render it more suitable for acquisition of symbiont-derived sterols.

### Phylogenetic Distribution of Animal SQase

Sulfoquinovosidase (SQase) is an enzyme that hydrolyzes sulfoquinovose (SQ) from sulfoquinovosyl diacylglycerol (SQDG) and from sulfoquinovosyl glycerol (SQGro), which is produced by deacylation of SQDG (Speciale et al. 2016). SQ is a sulfur-containing compound that is abundant in chloroplasts of photosynthetic organisms and is one of the most abundant organosulfur compounds on Earth (Fig. 6a) (Goddard-Borger & Williams 2017; Snow et al. 2021). Therefore, the putative SQase identified in the boring giant clam may contribute to host utilization of this abundant sulfur and lipid resource by degrading symbiont- and diet-derived SQDG.

**Fig. 6.**
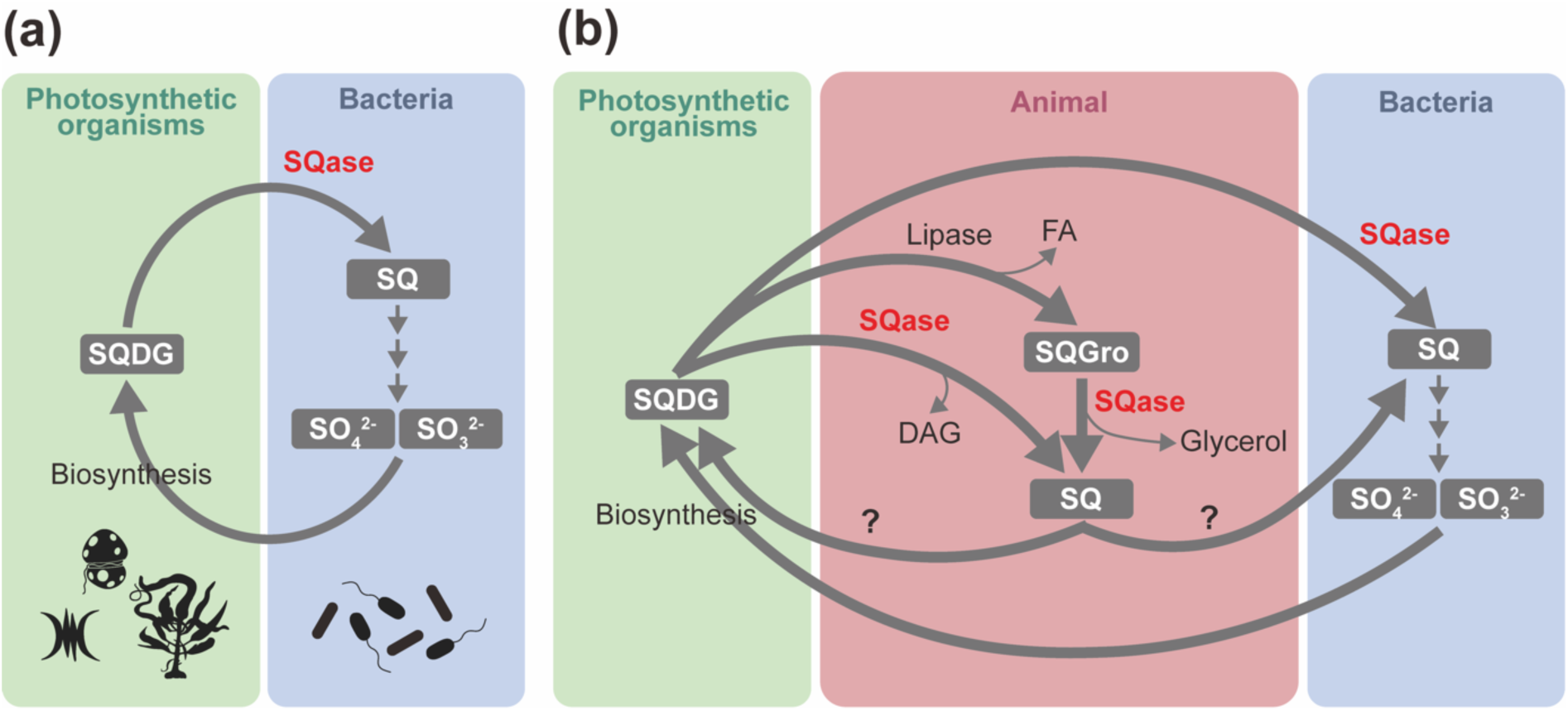
Roles of sulfoquinovosidase (SQase) in sulfoquinovose metabolism and sulfur cycling. **a)** Schematic overview of sulfur cycling mediated by sulfoquinovose (SQ) in the biosphere, modified from Goddard-Borger & Williams (2017) and Snow et al. (2021). Silhouettes of photosynthetic organisms and flagellated bacteria were obtained from PhyloPic. **b)** Conceptual model illustrating proposed roles of animal SQase in sulfur cycling. In photosymbiotic bivalves such as *Tridacna* and *Fragum*, symbiotic dinoflagellates correspond to photosynthetic organisms in this cycle, and SQ-mediated sulfur cycling may occur in the host body in association with associated bacterial communities. In this context, production and sharing of SQ may contribute to selective maintenance of SQ-degrading bacteria and potentially to recycling of SQ back to symbiotic dinoflagellates. SQ, sulfoquinovose; SQDG, sulfoquinovosyl diacylglycerol; SQGro, sulfoquinovosyl glycerol; SQase, sulfoquinovosidase; FA, fatty acid; DAG, diacylglycerol.

However, in annotation based on a BLAST search against the SwissProt database, the top hit for this protein was not an animal protein, but the SQase (YihQ) from *Escherichia coli* (strain K-12). Although Speciale et al. (2016) reported that some animal proteins show sequence similarity to bacterial SQase, the distribution of SQase homologs among animals and their phylogenetic relationships have not been examined in detail. Therefore, we first sought to determine whether animals possess SQase homologs. We searched for candidate genes in metazoan gene model datasets and public databases, and retained only those in which amino acids required for SQase activity and domain architecture were conserved, and then performed molecular phylogenetic analyses.

As a result, we identified genes that are inferred to be functional orthologs of bacterial SQases (*yihQ* from *E. coli* and *smoI* from *Agrobacterium fabrum*) (Speciale et al. 2016; Abayakoon et al. 2018) among the Arthropoda, Mollusca, Annelida, Brachiopoda, Bryozoa, Hemichordata, Echinodermata, Chordata, and Rotifera (hereafter referred to as “animal SQase”) (Figs. 2, 3, and 4). In the Arthropoda, animal SQase was detected only in Malacostraca, Copepoda, Thecostraca, Branchiopoda, Collembola (Hexapoda), and Ostracoda, and it was not detected in Insecta, the largest arthropod group. Among arthropods in which animal SQase was detected, only two species are terrestrial: *Armadillidium nasatum* (Malacostraca, Arthropoda; woodlouse) and *Orchesella cincta* (Hexapoda, Arthropoda; springtail). In Mollusca, animal SQase was detected in Bivalvia and Gastropoda. Similarly, in phyla other than Arthropoda and Mollusca, animal SQase was detected only in aquatic invertebrates. Moreover, even among aquatic invertebrates, animal SQase was not detected in generally carnivorous lineages, such as cephalopods or leeches. Taken together, these results suggest that animal SQase is widely distributed among aquatic invertebrates and may degrade dietary SQDG derived from algae, photosynthetic bacteria, or plants.

The gene tree of animal SQase broadly reflected the species phylogeny; however, only animal SQase sequences from bdelloid rotifers formed a clade with bacterial SQases, rather than with other animal SQases (Fig. 2). In addition, as described below, among animal phyla in which SQase homologs were detected, all rotifer species examined lacked signal peptides (Figs. 3 and 4b). These features raise the possibility that the SQase gene in rotifers was acquired via horizontal gene transfer (HGT) from bacteria. Indeed, bdelloid rotifers are particularly prone to HGT, and approximately 10% of their genes are thought to be derived from non-metazoan sources (Wilson et al. 2024).

### Molecular Features and Subcellular Localization of Animal SQase

In bacteria, SQase is thought to function in the cytosol after substrates are transported into the cell by membrane transporters (Snow et al. 2021). By contrast, if animal SQase is involved in digestion of dietary SQDG or SQGro, it would be expected to localize not in the cytosol, but in extracellular spaces such as the lumen of the digestive tract, or intracellular compartments such as lysosomes. To examine this possibility, we used machine-learning-based tools to predict signal peptides, which direct proteins to the secretory pathway in eukaryotes and mediate translocation across the plasma (inner) membrane in prokaryotes. As a result, signal peptides were predicted in many animals except rotifers, whereas they were not predicted in bacterial SQases (Figs. 3 and 4b).

We next predicted membrane anchoring via glycosylphosphatidylinositol (GPI) anchors and found that GPI-anchor attachment was predicted in many molluscs and in some echinoderms and chordates, indicating membrane association in these taxa (Figs. 3 and 4b). In addition, transmembrane helices were predicted in some echinoderms and chordates. Notably, the predicted GPI-anchor attachment sites (ω-sites) and transmembrane regions were located downstream (toward the C-terminus) of glycosyl hydrolase family 31 domains responsible for enzymatic activity (Fig. 4b). This topology suggests that in animal SQases possessing GPI-anchor attachment sites or transmembrane regions, the catalytic domain is exposed to the extracellular side and can act on extracellular substrates. Overall, these results suggest that animal SQases in these taxa enter the secretory pathway and are subsequently targeted to intracellular organelle lumens such as lysosomes, to the plasma membrane, or to the extracellular space, although their precise localization may differ among lineages. By contrast, no signal peptides were predicted for rotifer SQases (Fig. 3 and 4b), consistent with the idea that these genes were acquired via horizontal gene transfer from bacteria and have not subsequently acquired signal peptide sequences.

Because signal peptides are present in most animal SQases (Fig. 3 and 4b), they may have been acquired by the common ancestor of these animal lineages. Whereas GPI-anchoring was not predicted in annelids, brachiopods, or bryozoans, it was predicted in most molluscs, particularly bivalves, suggesting that GPI-anchor attachment may have been acquired in the ancestor of these lineages. In contrast, no GPI-anchor attachment was predicted in any species of giant clam, raising the possibility that in their common ancestor, animal SQase reverted to a secreted or lysosome-resident form. In addition, although sites inferred to be under positive selection in *Tridacna* and *Fragum* were located in or near the glycosyl hydrolase family 31 domain, amino acids critical for SQase activity were conserved (Fig. 4a). These observations suggest that substitutions fixed by positive selection may have altered properties such as optimal conditions without changing substrate specificity or catalytic activity.

### Physiological Roles of Animal SQase

To infer physiological roles of animal SQase in vivo, we examined tissue-specific gene expression patterns using publicly available RNA-seq datasets. In the bivalve *Mercenaria mercenaria*, the gastropod *Aplysia californica*, and the cephalochordate *Branchiostoma lanceolatum*, animal SQase showed significantly higher expression in digestive tissues such as the hepatopancreas (Fig. 5). These results suggest that across diverse animal lineages, animal SQase functions in catabolism of SQDG in the digestive tract. SQDG is widely distributed among phytoplankton, including diatoms, haptophytes, dinoflagellates, green algae, and cyanobacteria (Ma et al. 2025). As these organisms frequently constitute food sources for aquatic invertebrates, it is plausible that these animals possess SQDG-degrading enzymes, such as SQase, as part of their digestive enzyme repertoire.

In giant clams (*T. crocea* and *T. squamosa*), animal SQase was likewise highly expressed in the hepatopancreas, but in addition, it showed high expression in the outer mantle (Fig. 5). The outer mantle is the tissue primarily involved in harboring symbiotic dinoflagellates. Therefore, in giant clams, SQDG derived from symbiotic dinoflagellates may be degraded not only in digestive organs, but also directly in the tissue itself. Consistent with this interpretation, SQDG is abundant in both the outer mantle and the hepatopancreas (digestive diverticula) of *T. crocea*, indicating that substrates for animal SQase are readily available in these tissues (Sakai et al. 2023).

Histological observations suggested that in the outer mantle of giant clams, amoebocytes selectively engulf a subset of symbiotic dinoflagellates (Morton 1978), and that intracellular digestion by amoebocytes may occur in the hepatopancreas (Fankboner 1971). Based on these observations, animal SQase in giant clams may localize to lysosomes or related intracellular compartments in amoebocytes and participate in intracellular digestion in both the hepatopancreas and the outer mantle. In this scenario, evolutionary loss of GPI-anchor attachment may reflect a shift in the mode of digestion from extracellular to intracellular. Alternatively, animal SQase in giant clams may be secreted into extracellular spaces. Because zooxanthellal tubes distributed throughout the mantle extend from the stomach (Norton et al. 1992), they may represent a specialized structure derived from the digestive tract. Therefore, it would not be surprising if digestive enzymes, including SQase, were secreted into the lumen of these structures, as occurs in the digestive tract. Because zooxanthellal tubes represent a unique microenvironment not found in non-symbiotic bivalves, amino acid substitutions optimized for function in this specialized environment may have been fixed by positive selection in animal SQase of photosymbiotic bivalves.

### Integrated Perspectives on Animal SQase in Sulfur Cycling, Symbiosis, and Evolution

Sulfoquinovose (SQ) is one of the most abundant organic sulfur compounds on Earth, and biosynthesis and degradation of sulfoquinovosyl diacylglycerol (SQDG) is essential in global sulfur and carbon cycling, a process that is apparently driven primarily by bacteria (Fig. 6a) (Goddard-Borger & Williams 2017; Snow et al. 2021). In this study, however, we found that animal SQase genes are distributed in many aquatic invertebrate lineages. These findings suggest that in aquatic ecosystems, invertebrate animals may also contribute to sulfur and carbon cycling mediated by SQ.

As described above, animal SQase is expressed in digestive organs of aquatic invertebrates and is inferred to function as a digestive enzyme. Unlike bacterial SQases, animal SQases appear to have acquired signal peptides, enabling them to function in extracellular spaces or in intracellular vesicles. Furthermore, in certain lineages, additional evolutionary changes such as acquisition and subsequent loss of GPI-anchor attachment, acquisition of transmembrane domains, and amino acid substitutions driven by positive selection appear to have occurred. These changes likely represent adaptations to lineage-specific microenvironments and molecular functions in the animal body. In particular, in photosymbiotic giant clams, tissues with gene expression of animal SQase include the outer mantle, the primary symbiosis site, presumably to facilitate degradation of SQDG derived from symbiotic algae (Fig. 5).

In bacteria, SQ is further degraded and utilized through several metabolic pathways, depending on the species (Snow et al. 2021). In contrast, the presence of SQ-degrading gene clusters has not been reported in animals. Consistent with this, BLAST searches against the *T. crocea* genome and gene models failed to identify genes encoding the first enzyme of four known SQ catabolic pathways (Tables S3 and S4). These results suggest that the animal host is unlikely to directly utilize SQ generated through SQDG degradation. In mammals, few gut bacteria can degrade SQ, and dietary SQ is thought to influence the composition of the gut microbiota (Hanson et al. 2021; Krasenbrink et al. 2025). Similarly, in aquatic invertebrates such as giant clams, SQ degradation may be carried out by symbiotic microorganisms (Fig. 6b). Moreover, by producing SQ, the host may selectively maintain SQ-degrading bacteria in its organs. Giant clams harbor bacterial communities that differ markedly from those in surrounding seawater (Rossbach et al. 2019; Uchida, Li, et al. 2025), and SQ supply may represent one mechanism contributing to maintenance of these specialized microbiomes. It is also conceivable that SQ could be recycled to symbiotic dinoflagellates, and that the host regulates algal growth by controlling SQ availability (Fig. 6b). In cnidarians, it has been proposed that the supply of nutrients, including nitrogen, helps to regulate proliferation of symbiotic dinoflagellates (Cui et al. 2019; Xiang et al. 2020; Cui et al. 2022), and SQ may serve a comparable function in giant clams. Of course, it is also possible that the primary nutritional benefit of SQDG or SQGro degradation for the host lies in the diacylglycerol (DAG) or glycerol released during this process, utilizable lipid- and carbon-based energy sources for animals.

In this study, we identified SQase homologs in many animal lineages. However, because our searches primarily relied on gene and protein databases rather than complete genome sequences, SQase homologs may have been overlooked in species with incomplete gene prediction. More comprehensive genome-based searches will be necessary to accurately determine presence and absence SQase genes in each lineage. In addition, in vivo validation of protein subcellular localization will reveal SQase functions. Conserved and distinct tissue-specific expression patterns inferred from RNA-seq data should be corroborated by immunohistochemistry. Such future studies will further clarify contributions of animals to global sulfur and carbon cycling via SQ. They will also document evolutionary diversification of algal-derived metabolite utilization in invertebrates, and the role of host metabolism in shaping associated microbial communities.

## Materials and Methods

### Clustering of Orthologous Genes and Phylogenetic Analysis

Genomes and gene models of the following 12 bivalve species were used: *Magallana gigas* (*Crassostrea gigas*) (GCF_902806645.1), *Modiolus philippinarum* (Sun et al. 2017), *Mizuhopecten yessoensis* (GCF_002113885.1), *Ruditapes philippinarum* (GCF_026571515.1), *Tridacna crocea* (Uchida, Yamashita, et al. 2025), *Tridacna derasa* (GCA_963210305.1), *Tridacna gigas* (GCA_945859785.2), *Fragum fragum* (GCA_946902895.1), *Fragum whitleyi* (GCA_948146395.1), *Fragum sueziense* (GCA_963680895.1), *Cerastoderma edule* (GCA_963989375.1), and *Serripes groenlandicus* (GCA_031761405.1). For *T. crocea, T. derasa, T. gigas, F. fragum, F. whitleyi, and F. sueziense, C. edule, and S. groenlandicus*, gene models, predicted protein sequences, and functional annotations (BLASTP search against SwissProt and InterProScan search) reported in Uchida, Yamashita, et al. (2025) were reused. Results of orthogroup clustering performed with OrthoFinder in that study were also reused for the present analysis. In this OrthoFinder analysis, in addition to the 12 bivalve species, 10 metazoan species were included: two gastropods (*Lottia gigantea* and *Aplysia californica*), one cephalopod (*Octopus bimaculoides*), one arthropod (*Drosophila melanogaster*), two chordates (*Homo sapiens* and *Branchiostoma floridae*), and three cnidarians (*Exaiptasia diaphana*, *Nematostella vectensis*, and *Acropora tenuis*) (Uchida, Yamashita, et al. 2025).

Orthogroups shared by 12 bivalve species as single-copies were used for concatenated phylogenetic analysis. All amino acid sequences belonging to the same gene family were aligned using MAFFT v.7.310 with the ‘--auto’ option, and all gaps were removed using TrimAL v.1.4 with the ‘-nogaps’ option. All sequences from the same species were concatenated, and a maximum likelihood phylogenetic analysis was executed using RAxML v.8.2.12 with 100 bootstrap replicates and the ‘PROTOGAMMAAUTO’ option.

### Exploration of Lineage-Specific Positive Selection

To detect lineage-specific positive selection, we focused on 2,014 single-copy orthologous gene families shared among 12 bivalve species. Protein sequences were aligned using MAFFT v.7.310 with the -auto option (Katoh & Standley 2013), and codon alignments were generated using PAL2NAL v.14 (Suyama et al. 2006) with the -nogap option to remove codons corresponding to gapped amino-acid positions.

Screening for positive selection was conducted using ete-evol in the ETE toolkit v.3.1.3 (Huerta-Cepas et al. 2016), which automates execution of CODEML in the PAML package (Yang 2007) and subsequent likelihood ratio tests. The branch-site model was applied, and bsA1 (model = 2; NSsite = 2; fix_omega = 1; omega = 1) and bsA (model = 2; NSsite = 2; fix_omega = 0) models were used as null and alternative models, respectively. The species tree constructed as described above was used as the guide tree. The genera *Tridacna* and *Fragum* were analyzed independently as foreground lineages. Genes with a likelihood ratio test *p*-value < 0.05 were considered candidates for positive selection. Sites with BEB posterior probability > 0.95 were regarded as positively selected.

### Identification of Sulfoquinovosidase (SQase) Homologs

To identify sulfoquinovosidase (SQase) homologs in animals, we performed a three-step procedure as described below.

#### Step 1: Initial Identification of SQase-like Proteins

The *T. crocea* SQase-like protein was used as a query to search the nr and SwissProt databases maintained on the NIG supercomputer system using BLASTP (E-value < 10^-5^). From nr BLASTP hits, only animal-derived protein sequences were retained. In addition, (1) animal BLASTP hits against nr, (2) BLASTP hits against SwissProt, (3) proteins belonging to the orthogroup that includes the *T. crocea* SQase-like protein, and (4) putative SQase sequences reported by Speciale et al. (2016) were combined for phylogenetic analysis. Sequences were aligned using MAFFT with the --auto option, trimmed with TrimAL using the -gappyout option, and poorly aligned sequences were removed manually. A maximum likelihood phylogeny was inferred using RAxML with the -m PROTGAMMAAUTO option.

#### Step 2: Extraction of SQase Candidates Based on Phylogenetic Position

Based on topology of the Step 1 phylogenetic tree, putative SQase sequences were selected, because other proteins can cluster separately as non-SQase glucosidases of GH31 family. Proteins that formed a clade with *Escherichia coli* SQase (YihQ; SwissProt accession P32138) (Speciale et al. 2016) and *Agrobacterium* fabrum SQase (SmoI; SwissProt accession A9CEZ0) (Abayakoon et al. 2018) were retained and subjected to a second round of molecular phylogenetic analysis, which was performed using the same methods described above. To improve classification accuracy, known glucosidases downloaded from SwissProt were added as outgroup sequences.

#### Step 3: Functional Filtering and Refinement of Candidate Sequences

SQase candidates were further filtered using functional and structural criteria: (1) Sequences lacking or substituting several of the seven amino acids important for SQase catalytic activity (described below) were excluded because they may be pseudogenized or nonfunctional proteins. (2) When multiple isoforms were present for the same gene, the longest isoform was retained. (3) Candidates were required to possess a domain architecture consistent with a typical SQase, defined as a single Glycosyl hydrolase family 31 TIM-barrel domain (PF01055) as predicted by InterProScan. (4) The *F. whitleyi* sequence was manually curated to join fragmented predictions that likely resulted from gene model fragmentation. A final phylogenetic analysis was performed using the curated sequence set.

### Search for Downstream Genes of SQase in SQ Degradation Pathways

In bacteria, four distinct sulfoquinovose (SQ) degradation pathways have been described: the sulfo-EMP, sulfo-ED, sulfo-SFT, and sulfo-SMO pathways (Snow et al. 2021). Amino acid sequences of the first enzyme in each pathway (UniProt accessions: P32140, P0DOV5, A0A7W3RDA3, and A9CEY7) were retrieved from the UniProt/SwissProt database. These sequences were used as queries for TBLASTN searches against gene models and genome sequences of *Tridacna crocea*, with E-value < 1 × 10⁻³. For gene models or genomic regions showing significant similarity, functional annotations reported in Uchida, Yamashita, et al. (2025) were examined to assess whether homologs of known bacterial SQ degradation enzymes were present.

### Functional Inference of Animal SQase Homologs and Reassessment of Positive Selection

Amino acid sequences of candidate animal SQase homologs and representative bacterial SQases were aligned with MAFFT v7 using the --auto option. The presence of amino acids corresponding to binding sites and active sites of YihQ (*E. coli* SQase) reported by Speciale et al. (2016) was visually examined in resulting alignments. Signal peptides were predicted using SignalP 6.0 (Teufel et al. 2022), and glycosylphosphatidylinositol (GPI) anchor attachment sites were predicted using NetGPI 1.1 (Gíslason et al. 2021). Transmembrane helices were predicted using DeepTMHMM v1.0.44 (Hallgren et al. 2022).

For SQase sequences from the 12 bivalve species, amino acid regions corresponding to signal peptides and regions located downstream of the predicted GPI-anchor attachment sites (ω-sites) were removed. The remaining sequences, corresponding to the predicted mature proteins, were aligned using MAFFT with the --auto option. Poorly aligned regions were trimmed using trimAl with the -gappyout option. Molecular phylogenetic analyses were performed using RAxML with the PROTGAMMAAUTO model. Positive selection analyses by CODEML (Yang 2007) were conducted using ETE toolkit v.3.1.3 (Huerta-Cepas et al. 2016) following the same procedures described above.

### Comparative Analysis of Tissue-Specific SQase Gene Expression in Various Animals

To compare tissue-specific expression patterns of animal sulfoquinovosidase (SQase) genes, publicly available RNA-seq datasets and previously published transcriptomic studies were analyzed. For *Tridacna crocea*, tissue-specific RNA-seq data and corresponding quantification and statistical analysis results generated by Uchida, Yamashita, et al. (2025) were reused (n = 3 biological replicates).

For *Tridacna squamosa*, RNA-seq data were obtained from the NCBI Sequence Read Archive (SRA) under the accession numbers SRR13022154, SRR13022156–SRR13022165, SRR13022167–SRR13022176, and SRR13022178–SRR13022180 (n = 3 biological replicates). A previously reported de novo assembled transcriptome of *T. squamosa* (Uchida, Yamashita, et al. 2025) was used as the reference. RNA-seq reads were trimmed with TrimGalore v.0.6.10 (https://github.com/FelixKrueger/TrimGalore) and aligned to the reference transcriptome and quantified using the Trinity v.2.15.1 utility script, align_and_estimate_abundance.pl, with Bowtie2 employed for read alignment and RSEM for transcript-level abundance estimation (Haas et al. 2013). Transcript-level expected counts were used for downstream statistical analyses. Differential expression analysis among tissues was conducted using edgeR v.4.2.2 (Robinson et al. 2010) in R v.4.4.2. Minimally expressed transcripts were filtered out using the filterByExpr function prior to statistical testing. Library sizes were normalized using the trimmed mean of M-values (TMM) method. Differential expression was assessed using generalized linear models (GLMs) with quasi-likelihood F-tests. Resulting *p*-values were adjusted for multiple testing using the Benjamini-Hochberg procedure. Counts per million (CPM) and transcripts per million (TPM) values were also calculated.

For *Mercenaria mercenaria* and *Aplysia californica*, tissue-specific RNA-seq-based gene expression data expressed as TPM were retrieved from MolluscDB (Liu et al., 2025), with three biological replicates (n = 3). Differences in gene expression levels among tissues were assessed using one-way analysis of variance (ANOVA), followed by Tukey’s honestly significant difference (HSD) post hoc test, as implemented in R.

For *Branchiostoma lanceolatum*, tissue-specific RNA-seq data reported in a previous study were obtained from the NCBI Gene Expression Omnibus (GEO) under accession number GSE106430 (Marlétaz et al. 2018). Because only two biological replicates were available for this species (n = 2), no formal statistical testing was performed, and gene expression patterns were compared descriptively.

## Supporting information

Supplementary Figures S1 to S4 and Legends for Supplementary Tables S1 to S4

## Data Availability

No new data were generated or analysed in the course of this research.

## Acknowledgements

We thank Dr. Steven D. Aird for editing the manuscript. Computations were partially performed on the NIG supercomputer at ROIS National Institute of Genetics. This work was supported by Japan Society for the Promotion of Science (JSPS; grant number 21H04742, 24KJ0896, 24K01847, and 25H00948).

## Conflict of Interest

Authors declare that they have no competing interests.

## Notes

### Competing Interest Statement

The authors have declared no competing interest.

## References

Abayakoon P et al. 2018. Structural and biochemical insights into the function and evolution of sulfoquinovosidases. ACS Cent. Sci. 4:1266–1273.

Armstrong EJ, Roa JN, Stillman JH, Tresguerres M. 2018. Symbiont photosynthesis in giant clams is promoted by V-type H+-ATPase from host cells. J. Exp. Biol. 221. doi: 10.1242/jeb.177220.

Baker AC. 2003. Flexibility and Specificity in Coral-Algal Symbiosis: Diversity, Ecology, and Biogeography of Symbiodinium. Annu. Rev. Ecol. Evol. Syst. 34:661–689.

Blackall LL, Wilson B, Van Oppen MJH. 2015. Coral-the world’s most diverse symbiotic ecosystem. Mol. Ecol. 24:5330–5347.

Cui G et al. 2019. Host-dependent nitrogen recycling as a mechanism of symbiont control in Aiptasia. PLoS Genet. 15:e1008189.

Cui G et al. 2022. Nutritional control regulates symbiont proliferation and life history in coral-dinoflagellate symbiosis. BMC Biol. 20:103.

Dani V et al. 2017. Expression patterns of sterol transporters NPC1 and NPC2 in the cnidarian-dinoflagellate symbiosis. Cellular Microbiology. 19:e12753.

Dani V, Ganot P, Priouzeau F, Furla P, Sabourault C. 2014. Are Niemann-Pick type C proteins key players in cnidarian-dinoflagellate endosymbioses? Mol. Ecol. 23:4527–4540.

Dunn CW, Giribet G, Edgecombe GD, Hejnol A. 2014. Animal phylogeny and its evolutionary implications. Annu. Rev. Ecol. Evol. Syst. 45:371–395.

Fankboner PV. 1971. Intracellular digestion of symbiontic zooxanthellae by host amoebocytes in giant clams (Bivalvia: Tridacnidae), with a note on the nutritional role of the hypertrophied siphonal epidermis. Biol. Bull. 141:222–234.

Farmer MA, Fitt WK, Trench RK. 2001. Morphology of the symbiosis between Corculum cardissa (Mollusca: Bivalvia) and Symbiodinium corculorum (Dinophyceae). Biol. Bull. 200:336–343.

Gilbert SF, Sapp J, Tauber AI. 2012. A symbiotic view of life: we have never been individuals. Q. Rev. Biol. 87:325–341.

Giribet G, Edgecombe GD. 2019. The phylogeny and evolutionary history of arthropods. Curr. Biol. 29:R592–R602.

Gíslason MH, Nielsen H, Almagro Armenteros JJ, Johansen AR. 2021. Prediction of GPI-anchored proteins with pointer neural networks. Curr. Res. Biotechnol. 3:6–13.

Goddard-Borger ED, Williams SJ. 2017. Sulfoquinovose in the biosphere: occurrence, metabolism and functions. Biochem. J. 474:827–849.

Gupta SD, Sastry PS. 1987. Metabolism of the plant sulfolipid--sulfoquinovosyldiacylglycerol: degradation in animal tissues. Arch. Biochem. Biophys. 259:510–519.

Haas B et al. 2013. De novo transcript sequence reconstruction from RNA-seq using the Trinity platform for reference generation and analysis. Nat. Protoc. 8:1494–1512.

Hallgren J et al. 2022. DeepTMHMM predicts alpha and beta transmembrane proteins using deep neural networks. bioRxiv. doi: 10.1101/2022.04.08.487609.

Hambleton EA et al. 2019. Sterol transfer by atypical cholesterol-binding NPC2 proteins in coral-algal symbiosis. Elife. 8:1–26.

Hanson BT et al. 2021. Sulfoquinovose is a select nutrient of prominent bacteria and a source of hydrogen sulfide in the human gut. ISME J. 15:2779–2791.

Huerta-Cepas J, Serra F, Bork P. 2016. ETE 3: Reconstruction, Analysis, and Visualization of Phylogenomic Data. Mol. Biol. Evol. 33:1635–1638.

Iguchi A, Shinzato C, Forêt S, Miller DJ. 2011. Identification of fast-evolving genes in the scleractinian coral Acropora using comparative EST analysis. PLoS One. 6:e20140.

Ip YK, Chew SF. 2021. Light-Dependent Phenomena and Related Molecular Mechanisms in Giant Clam-Dinoflagellate Associations: A Review. Frontiers in Marine Science. 8:1–23.

Ishii Y et al. 2025. Positive selection of a starch synthesis gene and phenotypic differentiation of starch accumulation in symbiotic and free-living coral symbiont dinoflagellate species. Genome Biol. Evol. 17:evaf133.

Katoh K, Standley DM. 2013. MAFFT Multiple Sequence Alignment Software Version 7: Improvements in Performance and Usability. Mol. Biol. Evol. 30:772–780.

Kirkendale L. 2009. Their Day in the Sun: molecular phylogenetics and origin of photosymbiosis in the ‘other’ group of photosymbiotic marine bivalves (Cardiidae: Fraginae). Biological Journal of The Linnean Society. 97:448–465.

Krasenbrink J et al. 2025. Sulfoquinovose is exclusively metabolized by the gut microbiota and degraded differently in mice and humans. Microbiome. 13:184.

Levy S et al. 2021. A stony coral cell atlas illuminates the molecular and cellular basis of coral symbiosis, calcification, and immunity. Cell. 184:2973–2987.e18.

Li J et al. 2020. Shedding light: A phylotranscriptomic perspective illuminates the origin of photosymbiosis in marine bivalves. BMC Evol. Biol. 20:1–15.

Li J, Volsteadt M, Kirkendale L, Cavanaugh CM. 2018. Characterizing Photosymbiosis Between Fraginae Bivalves and Symbiodinium Using Phylogenetics and Stable Isotopes. Frontiers in Ecology and Evolution. 6. doi: 10.3389/fevo.2018.00045.

Li Jun et al. 2024. Chromosome-level genome assembly and annotation of rare and endangered tropical bivalve, *Tridacna crocea*. Scientific Data. 11:1–10.

Li R, Leiva C, Lemer S, Kirkendale L, Li J. 2025. Photosymbiosis shaped animal genome architecture and gene evolution as revealed in giant clams. Commun. Biol. 8:7.

Li Ruiqi, Zarate D, Avila-Magaña V, Li J. 2024. Comparative transcriptomics revealed parallel evolution and innovation of photosymbiosis molecular mechanisms in a marine bivalve. Proc. Biol. Sci. 291:20232408.

Liu H et al. 2018. Symbiodinium genomes reveal adaptive evolution of functions related to coral-dinoflagellate symbiosis. Commun. Biol. 1:95.

Ma X et al. 2025. Cosmopolitan marine bacteria facilitate a vast phytoplankton-derived sulfonate-based carbon flow through sulfoquinovosidases. Nat. Commun. 17:209.

Marlétaz F et al. 2018. Amphioxus functional genomics and the origins of vertebrate gene regulation. Nature. 564:64–70.

McFall-Ngai M et al. 2013. Animals in a bacterial world, a new imperative for the life sciences. Proc. Natl. Acad. Sci. U. S. A. 110:3229–3236.

McKenna V et al. 2021. The Aquatic Symbiosis Genomics Project: probing the evolution of symbiosis across the Tree of Life. Wellcome Open Res. 6:254.

Melo Clavijo J, Donath A, Serôdio J, Christa G. 2018. Polymorphic adaptations in metazoans to establish and maintain photosymbioses. Biol. Rev. Camb. Philos. Soc. 93:2006–2020.

Mies M. 2019. Evolution, diversity, distribution and the endangered future of the giant clam–Symbiodiniaceae association. Coral Reefs. 38:1067–1084.

Mies M, Sumida PYG, Rädecker N, Voolstra CR. 2017. Marine invertebrate larvae associated with Symbiodinium: A mutualism from the start? Frontiers in Ecology and Evolution. 5:1–11.

Morishima SY et al. 2019. Study on expelled but viable zooxanthellae from giant clams, with an emphasis on their potential as subsequent symbiont sources. PLoS One. 14:1–20.

Morton B. 1978. The diurnal rhythm and the processes of feeding and digestion in Tridacna crocea (BivaMa: Tridacnidae). J. Zool. (1987). 185:371–387.

Norton JH, Shepherd MA, Long HM, Fitt WK. 1992. The zooxanthellal tubular system in the giant clam. Biol. Bull. 183:503–506.

Rahman IA, Zamora S. 2024. Origin and early evolution of echinoderms. Annu. Rev. Earth Planet. Sci. 52:295–320.

Robinson MD, McCarthy DJ, Smyth GK. 2010. edgeR: a Bioconductor package for differential expression analysis of digital gene expression data. Bioinformatics. 26:139–140.

Rossbach S, Cardenas A, Perna G, Duarte CM, Voolstra CR. 2019. Tissue-Specific Microbiomes of the Red Sea Giant Clam Tridacna maxima Highlight Differential Abundance of Endozoicomonadaceae. Front. Microbiol. 10. doi: 10.3389/fmicb.2019.02661.

Roth MS. 2014. The engine of the reef: photobiology of the coralâ€“algal symbiosis. Front. Microbiol. 5:422.

Sakai R et al. 2023. Smart utilization of betaine lipids in the giant clam Tridacna crocea. iScience. 26:107250.

Shikina S et al. 2024. Genome and tissue-specific transcriptomes of the large-polyp coral, Fimbriaphyllia (Euphyllia) ancora: a recipe for a coral polyp. Commun. Biol. 7:899.

Shinzato C et al. 2021. Eighteen Coral Genomes Reveal the Evolutionary Origin of Acropora Strategies to Accommodate Environmental Changes. Mol. Biol. Evol. 38:16–30.

Snow AJD, Burchill L, Sharma M, Davies GJ, Williams SJ. 2021. Sulfoglycolysis: catabolic pathways for metabolism of sulfoquinovose. Chem. Soc. Rev. 50:13628–13645.

Soo P, Todd PA. 2014. The behaviour of giant clams (Bivalvia: Cardiidae: Tridacninae). Mar. Biol. 161:2699–2717.

Speciale G, Jin Y, Davies GJ, Williams SJ, Goddard-Borger ED. 2016. YihQ is a sulfoquinovosidase that cleaves sulfoquinovosyl diacylglyceride sulfolipids. Nat. Chem. Biol. 12:215–217.

Subramanian K, Balch WE. 2008. NPC1/NPC2 function as a tag team duo to mobilize cholesterol. Proc. Natl. Acad. Sci. U. S. A. 105:15223–15224.

Sun J et al. 2017. Adaptation to deep-sea chemosynthetic environments as revealed by mussel genomes. 1:1–7.

Suyama M, Torrents D, Bork P. 2006. PAL2NAL: robust conversion of protein sequence alignments into the corresponding codon alignments. Nucleic Acids Res. 34:W609–12.

Teufel F et al. 2022. SignalP 6.0 predicts all five types of signal peptides using protein language models. Nat. Biotechnol. 40:1023–1025.

Uchida T, Yamashita H, et al. 2025. Genomic insights into photosymbiosis in giant clams: Comparisons with coral strategies. bioRxiv. 2025.11. 20.689417. doi: 10.1101/2025.11.20.689417.

Uchida T, Li Y, Yamashita H, Shimada G, Shinzato C. 2025. Microbiome of the boring giant clam provides insights into holobiont resilience under coral reef environmental stress. Environ. Microbiol. 27:e70161.

Voolstra CR et al. 2011. Rapid evolution of coral proteins responsible for interaction with the environment. PLoS One. 6:e20392.

Wilson CG, Pieszko T, Nowell RW, Barraclough TG. 2024. Recombination in bdelloid rotifer genomes: asexuality, transfer and stress. Trends Genet. 40:422–436.

Xiang T et al. 2020. Symbiont population control by host-symbiont metabolic interaction in Symbiodiniaceae-cnidarian associations. Nat. Commun. 11:108.

Yang Z. 2007. PAML 4: phylogenetic analysis by maximum likelihood. Mol. Biol. Evol. 24:1586–1591.

Yellowlees D, Rees TAV, Leggat W. 2008. Metabolic interactions between algal symbionts and invertebrate hosts. Plant Cell Environ. 31:679–694.

Yoshioka Y et al. 2023. Genes possibly related to symbiosis in early life stages of Acropora tenuis inoculated with Symbiodinium microadriaticum. Communications Biology. 6:1027.

Yuyama I, Ishikawa M, Nozawa M, Yoshida M-A, Ikeo K. 2018. Transcriptomic changes with increasing algal symbiont reveal the detailed process underlying establishment of coral-algal symbiosis. Sci. Rep. 8:16802.

