## Supplementary Figures S1 to S4 and Legends for Supplementary Tables S1 to S4 for "Possible Novel Sulfolipid Utilization Pathway in Giant Clams and Other Aquatic Invertebrates: Implications for Photosymbiosis and Sulfur Cycling"

Taiga Uchida *et al.*

### Legends for Supplementary Tables S1 to S4

**Table S1.**

Results of screening for positive selection in the *Tridacna* lineage. Orthogroups with *p*-values < 0.05 are shown.

**Table S2.**

Results of screening for positive selection in the *Fragum* lineage. Orthogroups with *p*-values < 0.05 are shown.

**Table S3.**

Results of screening for genes downstream of SQase in SQ degradation pathways against *T. crocea* gene models. TBLASTN results and annotations of the BLAST hits are shown. Human orthologs are based on OrthoFinder results.

**Table S4.**

Results of screening for genes downstream of SQase in SQ degradation pathways against the *T. crocea* genome. TBLASTN results are shown. All BLAST hit regions corresponded to parts of protein-coding genes; therefore, annotations of the corresponding genes are also provided. Human orthologs are based on OrthoFinder results.

### Supplementary Figures

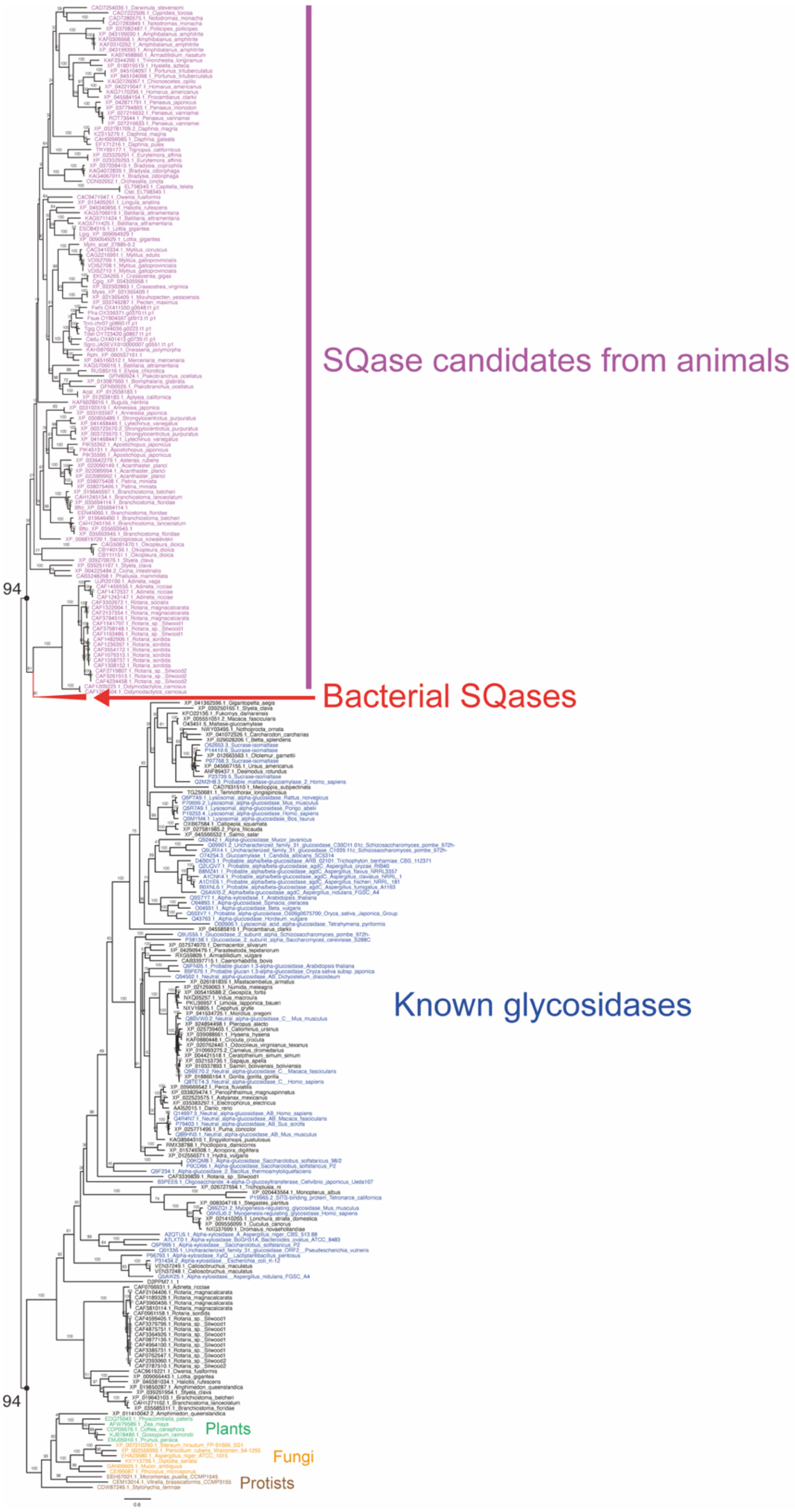

**Fig. S1.** Results of step 1 of animal SQase identification. Bootstrap values are shown on the branches.

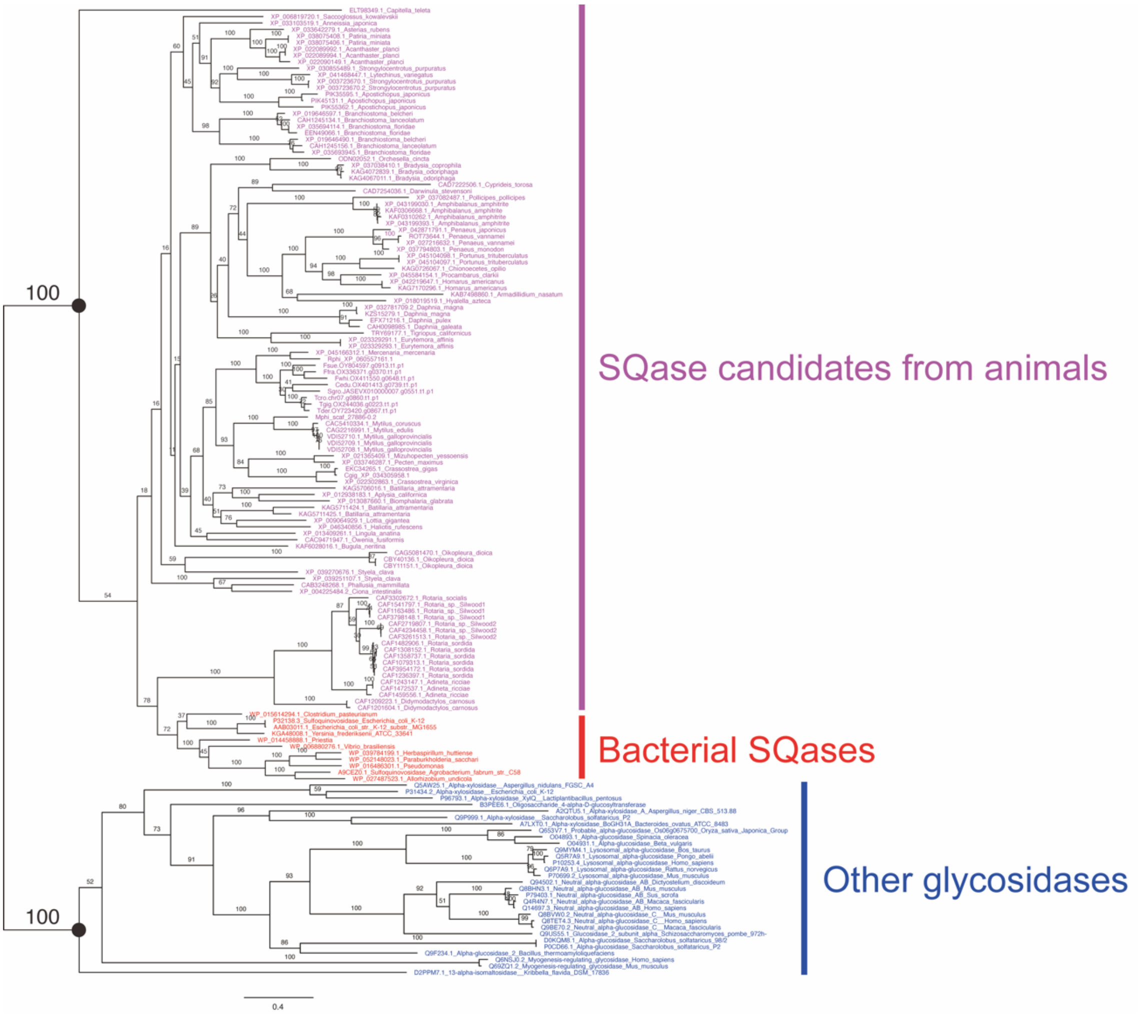


**Fig. S2.** Results of step 2 of animal SQase identification. Bootstrap values are shown on the branches.

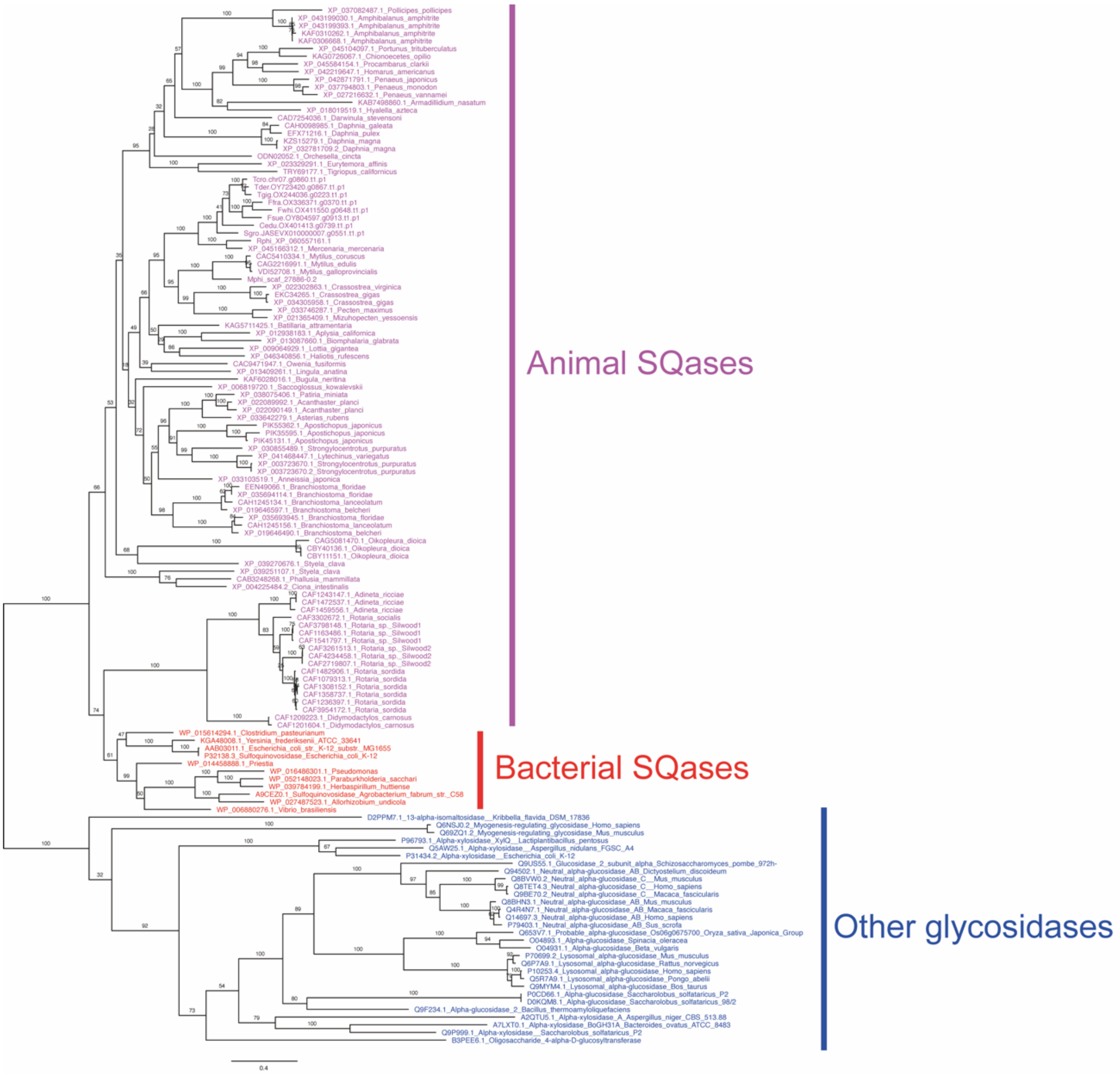


**Fig. S3.** Results of step 3 of animal SQase identification (full results corresponding to Fig. 2). Bootstrap values are shown on the branches.

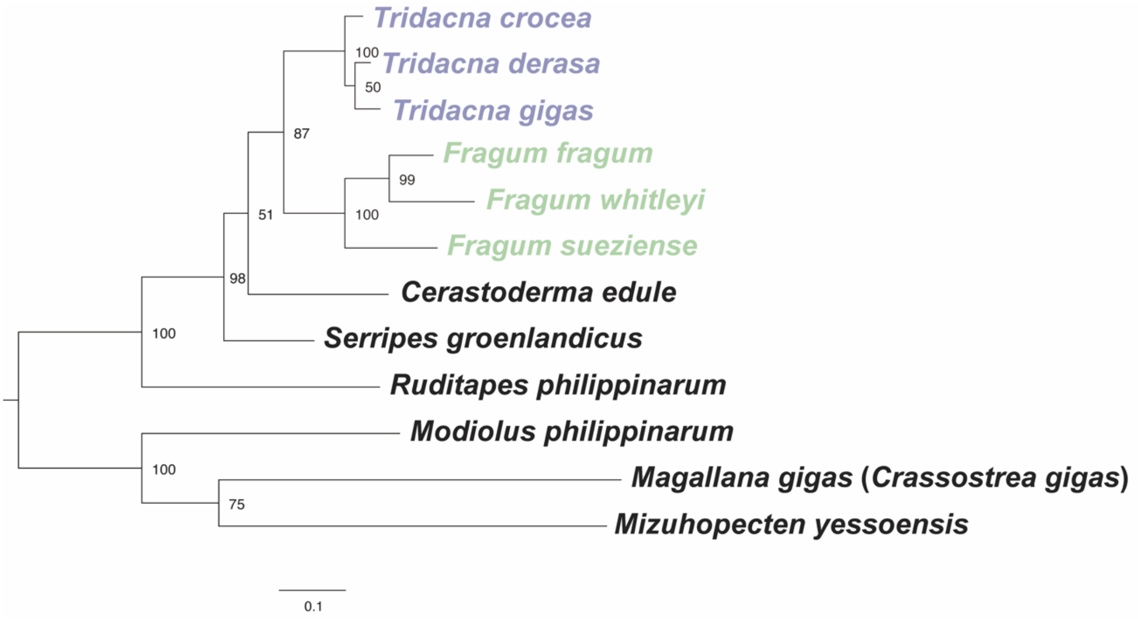


**Fig. S4.** Maximum likelihood molecular phylogenetic tree of animal SQases of 12 bivalve species used for detecting positive selection. Bootstrap values are shown on the nodes.
